# Fission yeast TRP channel Pkd2p localizes to the cleavage furrow and regulates cell separation during cytokinesis

**DOI:** 10.1101/316380

**Authors:** Zachary Morris, Debatrayee Sinha, Abhishek Poddar, Brittni Morris, Qian Chen

## Abstract

Force plays a central role in separating daughter cells during cytokinesis, the last stage of cell division. However, the mechanism of force-sensing during cytokinesis remains unknown. Here we discovered that Pkd2p, a putative force-sensing TRP channel, localizes to the cleavage furrow during cytokinesis of the fission yeast, *Schizosaccharomyces pombe*. Pkd2p, whose human homologues are associated with Autosomal Polycystic Kidney Disease, is an essential protein whose localization depends on the contractile ring and the secretory pathway. We identified and characterized a novel *pkd2* mutant *pkd2-81KD*. The *pkd2* mutant cells show signs of osmotic stress, including temporary shrinking, paused turnover of the cytoskeletal structures and hyper-activated MAPK signaling. During cytokinesis, although the contractile ring constricts more rapidly in the *pkd2* mutant than the wild-type cells (50% higher), the cell separation in the mutant is slower and often incomplete. These cytokinesis defects are also consistent with mis-regulated turgor pressure. Lastly, the *pkd2* mutant exhibits strong genetic interactions with two mutants of the SIN pathway, a signaling cascade essential for cytokinesis. We propose that Pkd2p modulates osmotic homeostasis and is potentially a novel regulator of cytokinesis.

**Highlight summary for TOC:** Fission yeast TRP channel Pkd2p is the homologue of human polycystins. The *pkd2* mutant exhibits defects in the contractile ring closure and cell separation during cytokinesis. This essential protein localizes to the cleavage furrow where it likely regulates osmotic homeostasis during cytokinesis.

## Introduction

Force plays a critical role in separating daughter cells during cytokinesis, the last stage of cell division (for review see (Pollard, 2010; Srivastava *et al.*, 2016)). Since the initial discovery (Rappaport, 1967), studies by many groups have established that the actomyosin contractile ring is essential to generate the force required for cytokinesis (De Lozanne and Spudich, 1987; Sanger and Sanger, 1980; Straight *et al.*, 2003). What remains largely unknown is whether and how dividing cells respond to this force, a potential mechanical stimulus. Some have proposed that the activity of myosin II, a motor protein in the ring, may be force-sensitive but the mechanism is far from clear (Effler *et al.*, 2006; Pinheiro *et al.*, 2017). On the other hand, few studies have examined whether any other mechanisms could contribute to force-sensing during cytokinesis.

Mechanosensitive (MS) channels can open and close in response to mechanical force (for review, see (Kung, 2005; Ranade *et al.*, 2015)). The first such channel was identified in bacteria (Martinac *et al.*, 1987). Since then, many more MS channels have been found in not only prokaryotes but also eukaryotes including humans. Among them, the best studied examples are *E.coli* MscS and MscL which modulate intracellular osmolarity in response to hypoosmotic shocks (Sukharev *et al.*, 1994; Sukharev *et al.*, 1996). Comparably, our understanding of eukaryotic MS channels such as PKD2, Piezo, and NompC is relatively limited, even though they are essential in sensing various forms of mechanical stimuli including flow, touch and osmolarity (Coste *et al.*, 2010; Mochizuki *et al.*, 1996; Walker *et al.*, 2000). In particular, we have not completely understood how MS channels sense force, but two widely accepted models have been proposed (Arnadottir and Chalfie, 2010; Kung, 2005; Ranade *et al.*, 2015). The “Membrane tension” model hypothesizes that force directly stretches the membrane where MS channels are located to open their ion pores. The “Tethered” model proposes that force is applied to the extracellular matrix and/or the intracellular cytoskeleton to activate the MS channels indirectly. The functions of MS channels in non-sensory cells also remain largely unexplored.

We propose to examine the potential roles of MS channels during cytokinesis using the fission yeast *Schizosaccharomyces pombe.* Classical genetics studies have identified a large number of important cytokinesis genes in this model organism (Balasubramanian *et al.*, 1998; Johnson *et al.*, 2012; Pollard and Wu, 2010). The molecular mechanism of fission yeast cytokinesis has been well studied and found to be largely conserved in higher eukaryotes (Pollard and Wu, 2010). Like animal cells, fission yeast assemble an actomyosin contractile ring at the cell division plane at the beginning of cytokinesis (Wu *et al.*, 2003). This process requires many essential proteins including Myo2p, Cdc12p and Cofilin (Balasubramanian *et al.*, 1998; Chang *et al.*, 1997; Chen and Pollard, 2011) as well as continuous polymerization of the actin filaments (Courtemanche *et al.*, 2016). The ring assembly is followed by the contractile ring closure driven by the activities of both type II and type V myosins (Laplante *et al.*, 2015; Mishra *et al.*, 2013). The final step of cytokinesis is cell separation (for review see (Garcia Cortes *et al.*, 2016; Sipiczki, 2007)), that includes many key steps such as the septum biosynthesis (Cortes *et al.*, 2002; Cortes *et al.*, 2012; Liu *et al.*, 1999; Munoz *et al.*, 2013) and the cell wall degradation (Dekker *et al.*, 2004; Martin-Cuadrado *et al.*, 2003).

Mechanical forces, including that from the contractile ring, the septum and the turgor, all play important roles during fission yeast cytokinesis. The tension originated from the contractile ring is required for the initiation of cell separation (Balasubramanian *et al.*, 1998; Mishra *et al.*, 2013; Stachowiak *et al.*, 2014). Just as importantly, the ring provides the mechanical cue that guides the septum biosynthesis (Thiyagarajan *et al.*, 2015; Zhou *et al.*, 2015). However, a recent work has shown that the ring is not required continuously during cytokinesis (Proctor *et al.*, 2012). The compression applied by the expanding septum helps both drive the cleavage furrow ingression and anchor the furrow at the cell division plane (Arasada and Pollard, 2014; Thiyagarajan *et al.*, 2015). This was vividly demonstrated by the observation that a defect in the septum biosynthesis often results in failed cell separation as well as post-separation cell lysis (Cortes *et al.*, 2012; Munoz *et al.*, 2013). Lastly, the turgor pressure, essential to morphogenesis of yeast cells, plays a key but often under-appreciated role in separating daughter cells (Abenza *et al.*, 2015; Atilgan *et al.*, 2015; Proctor *et al.*, 2012). Despite the importance of these forces, we have much to learn about how these mechanical stimuli are sensed and modulated during fission yeast cytokinesis, including the potential role of MS channels.

Here we examined the role of a putative transient receptor potential (TRP) channel Pkd2p during fission yeast cytokinesis. Many eukaryotic MS channels including NompC and human PKD2 belong to the TRP channel family (for review see (Christensen and Corey, 2007)). Consisting of seven subfamilies, this family of non-selective cation channels has been found in most eukaryotes (Wu *et al.*, 2010), including yeasts (Ma *et al.*, 2011; Palmer *et al.*, 2005; Palmer *et al.*, 2001). We discovered that a fission yeast TRP channel Pkd2p localizes to the cleavage furrow during cytokinesis. The *pkd2-81KD* mutation led to strong cytokinesis defects. Our genetic studies also identified the potential interaction between *pkd2* and the SIN pathway, a Hippo-like signaling cascade that regulates fission yeast cytokinesis (Hergovich and Hemmings, 2012; McCollum and Gould, 2001).

## Results

### A putative TRP channel Pkd2p localizes to the cleavage furrow

To identify MS channels that may play a role during cytokinesis, we first determined whether any of the fission yeast TRP channels localizes to the cell division plane (Aydar and Palmer, 2009; Ma *et al.*, 2011; Palmer *et al.*, 2005). The fission yeast genome encodes for the genes of three putative TRP channels, Pkd2p, Trp663p and Trp1322p (Pombase). We tagged each of the endogenous proteins with GFP at its C-terminus (unless specified, mEGFP was used throughout this study). Only Pkd2p-GFP localized to the cell division plane prominently during cytokinesis (Fig. 1A and S1A, Movie S1). The *pkd2::pkd2-GFP* cells exhibited similar morphology and viability to wild-type cells (data not shown), indicating that Pkd2p-GFP is a functional replacement of the endogenous protein. We concluded that Pkd2p is a putative TRP channel localized at the cleavage furrow and it may have a role in cytokinesis.

**Figure 1:**
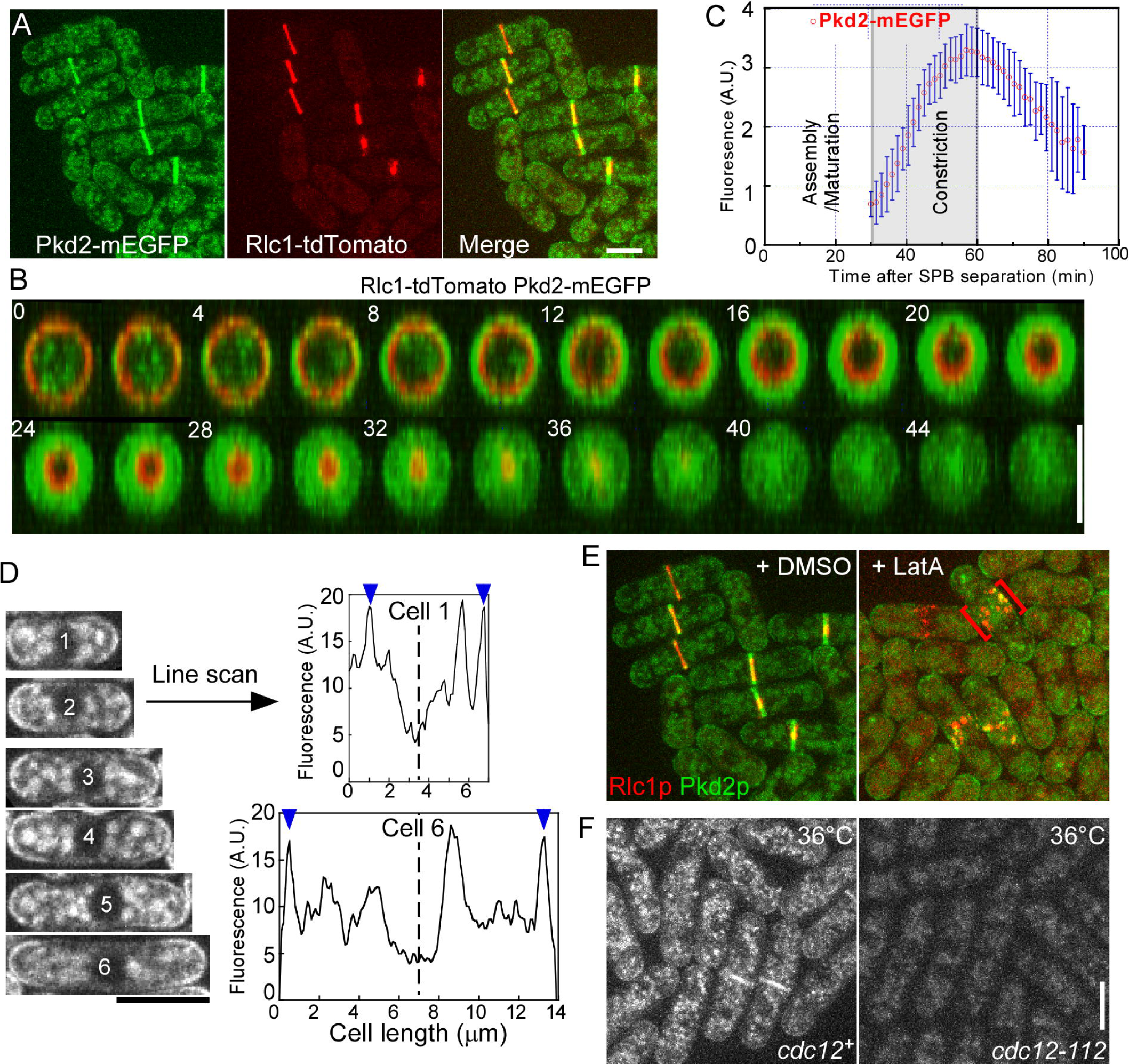
Localization of a TRP channel Pkd2p at the cell division plane. (A-C) Pkd2p localization during cytokinesis. (A-B) Fluorescence micrograph of cells expressing Pkd2p-GFP (green) and Rlc1p-tdTomoato (red), a marker for the contractile ring. Unless specified, maximum intensity projections are shown in all figures. (A) Fluorescence micrographs showing Pkd2p and Rlc1p co-localized to the contractile rings. Right: merged image. (B) Time-lapse micrographs of the division plane of a cell (Head-on view and merged color). Numbers represent time in mins. (C) A plot showing the time course of Pkd2p-GFP fluorescence at the cell division plane after the separation of SPBs (time zero). Pkd2p-GFP appeared at the division plane at the start of ring constriction (shaded area) and the fluorescence intensities peaked when the ring closure was completed. (D) Pkd2p localization during interphase. Left: Fluorescence micrographs of six cells expressing Pkd2p-GFP (numbered from 1-6 based on their length). Average intensity projections of three center Z-slices are shown. Right: Line scans based on the micrographs of cell 1 (top) and 6 (bottom). Pkd2p localized equally to the two cell tips (blue arrowheads). Dashed lines: median plane of the cells. (E-F) Regulation of Pkd2p localization. (E) Fluorescence micrographs of cells expressing Pkd2p-GFP, treated for one hour with either control (DMSO, left) or 10 µM latrunculin A (LatA, right). Disassembly of the contractile ring displaced Pkd2p-GFP to the cortex clumps (red brackets). (F) Fluorescence micrographs of wild-type (*cdc12*^*+*^) and *cdc12-112* cells expressing Pkd2p-GFP at 36°C. Pkd2p was displaced from the division plane at the restrictive temperature. Bars represent 5 µm. Error bars represent standard deviations (S.D.).

We determined Pkd2p localization throughout cell cycle using live fluorescence microscopy. During cell division, Pkd2p-GFP first appeared at the cell division plane during telophase, ∼30 mins after separation of the spindle pole bodies (SPB) (Fig. 1B and 1C). Its molecular number gradually increased, eventually peaking at ∼1,200, when the ring closure was completed (Fig. S1C). Pkd2p-GFP localized to the intracellular vesicles and organelles. It was also found at the plasma membrane of cell tips (Fig. 1D and Movie S1) but it didn’t show a preference for either one of the tips (Fig. 1D). We concluded that Pkd2p localization at the cell division plane is dependent upon cell-cycle progression.

We next examined how Pkd2p is recruited to the cleavage furrow during cytokinesis. First, we determined whether the actin or microtubule cytoskeletal structures are directly required for Pkd2p localization during cytokinesis. The actin cytoskeletal structures were disassembled within 5 mins in the cells treated with 50 µM Latrunculin A (Chen and Pollard, 2013; Coue *et al.*, 1987). However, Pkd2p-GFP remained at the cell division plane in these cells (Fig. S1D). Similarly, after the microtubules had been depolymerized in the cells treated by MBC (Sawin and Snaith, 2004), Pkd2p localization at the division plane was unchanged (Fig. S1E). These observations demonstrated that neither actin filaments nor microtubules are required to maintain Pkd2p localization at the cleavage furrow. Next, we determined whether the contractile ring was required for Pkd2p localization. We dissembled the contractile ring by treating the cells with 10 µM Latrunculin A for an extended period of time. In these cells, Pkd2p-GFP dispersed from the cleavage furrow to discrete puncta on the plasma membrane (Fig. 1E). Consistent with this finding, Pkd2p also failed to localize to the cleavage furrow in *cdc12-112*, a mutant of the formin required for the ring assembly (Chang *et al.*, 1997), at the restrictive temperature (Fig. 1F). Lastly, we determined whether the secretory pathway is required for Pkd2p localization. Brefeldin A (BFA) was used to inhibit the secretory pathway (Misumi *et al.*, 1986). In these BFA treated cells, Pkd2p-GFP localization to the cell division plane was inhibited (Fig. S1F). Together, our data suggested that Pkd2p localization at the cleavage furrow during cytokinesis requires both the contractile ring and the secretory pathway.

### *Pkd2* is an essential gene required for both cell growth and cell division

Pkd2p is the only fission yeast homologue of polycystins (Palmer *et al.*, 2005), an evolutionally conserved subfamily (TRPP) of TRP channels (Wu *et al.*, 2010). There are two human homologues of Pkd2p, PKD1 and the MS channel PKD2 (Hanaoka *et al.*, 2000; Nauli *et al.*, 2003). Mutations of either one of the human genes lead to a hereditary renal disorder Autosomal Polycystic Kidney Disease (ADPKD). As a putative TRP channel, fission yeast Pkd2p possesses a N-terminal signal peptide, an extracellular lipid binding (ML-like) domain, a central transmembrane domain (TRP domain) and a C-terminal cytoplasmic coiled-coil domain (Fig. 2A and 2B). Pkd2p had been proposed to be a part of the cell wall integrity pathway (Aydar and Palmer, 2009; Palmer *et al.*, 2005) but its role in cytokinesis had not been characterized.

**Figure 2:**
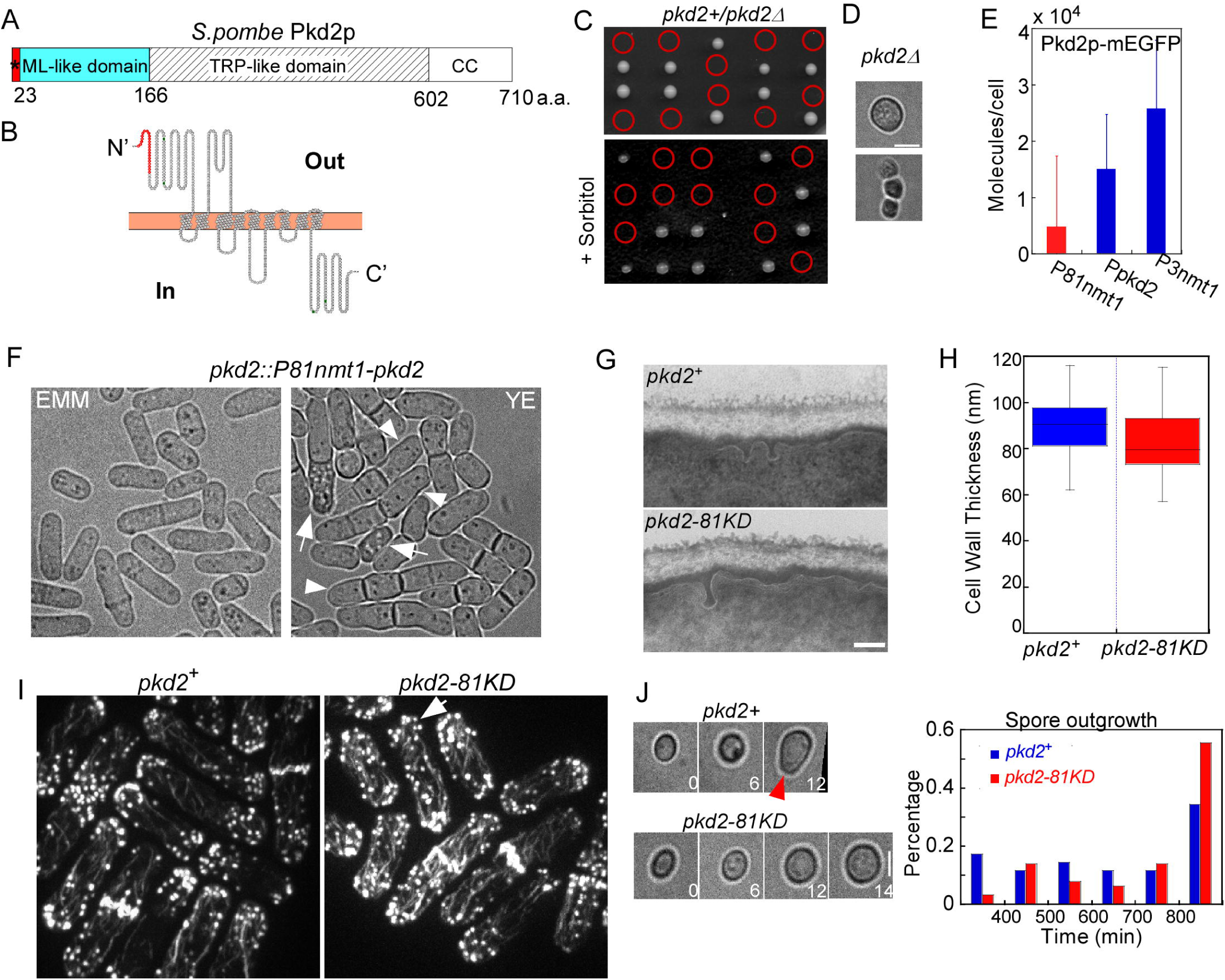
A essential gene *pkd2* is required for cell growth and cell division. (A-B) A putative TRP channel Pkd2p. (A) Domain structure of Pkd2p. *: the signal peptide. ML-like: MD-2-related lipid recognition domain. TRP-like: the transmembrane domain. CC: coiled-coil domain. (B) Predicted topology of Pkd2p on the plasma membrane (Orange). Red: the signal peptide. Pkd2p is projected to contain extensive extracellular and cytoplasmic domains. (C-D) *pkd2* is an essential gene. (C) Tetrad dissections of sporulated *pkd2*^*+*^*/pkd2Δ* cells, germinated on either YE5s (top) or YE5s plus 1.2 M sorbitol plates (bottom). Red circles: expected *pkd2Δ* spores. The *pkd2* deletion mutant is not viable. (D) Representative micrographs of *pkd2Δ* spores germinated on YE5s for more than one week. Most mutant spores died during the outgrowth (top) although a small number died after 1-2 rounds of cell division (bottom). Compared to wild-type spores that initiated polarized outgrowth (Fig. 2J), *pkd2Δ* spores grew isotropically. (E-J) Characterization of a hypomorphic *pkd2* mutant. (E) Bar graph comparing the average number of Pkd2p-GFP molecules per cell among *pkd2::P81nmt1-pkd2-GFP* (P81nmt1), *pkd2-GFP* (Ppkd2) and *pkd2::P3nmt1-pkd2-GFP* (P3nmt1) in YE5s (n > 60). Replacing the endogenous *pkd2* promoter with P81nmt1 reduced the number of Pkd2p molecules by 70%, while P3nmt1 promoter increased the molecular number one-fold. (F) Micrographs of *pkd2-81KD* cells in either YE5s (suppressing) or EMM5s (inducing) media. Under suppressing condition, many mutant cells were multi-septated (arrowheads) and some appeared to be lysed (arrows). Both morphological defects were rescued under inducing condition. (G) Transmission electron micrographs of the cell wall of a wild-type (top) and a *pkd2-81KD* (bottom) cells. (H) Box plot showing the average thickness of wild-type and *pkd2-81KD* cell wall (n > 8 cells). There was no significant difference between them (P > 0.05). (I) Fluorescence micrographs of Bodipy-Phallacidin stained wild-type (left) and *pkd2-81KD* (right) cells. Distribution of the actin patches (arrow) was less polarized in the mutant cells, even though their actin cytoskeletal structures appeared to be similar to those of the wild-type cells. (J) Left: Representative time-lapseb micrographs of a wild-type (top) and a *pkd2-81KD* (bottom) spores after germination (time zero). Numbers represent time in hours. Right: Histogram showing the elapsed times before the initiation of polarized outgrowth (arrowhead). The outgrowth of *pkd2-81KD* mutant (red) was delayed, compared to the wild-type spores (blue) (n > 30). Error bars represent standard deviations.

To determine the function of Pkd2p, we first attempted to construct a deletion mutant. However, *pkd2Δ* was not viable (Fig. 2C), which agreed with the previous finding (Palmer *et al.*, 2005). In fact, microscopic examination of *pkd2Δ* spores found that the majority of them (24/26) failed to divide and died before initiating polarized outgrowth (Fig. 2D), although few of the mutant spores (2/26) did divide 1-2 rounds before death (Fig. 2D). We attempted to rescue the deletion mutant with an osmotic stabilizer because of its proposed role in the cell wall integrity pathway. However, *pkd2Δ* did not survive in the presence of 1.2 M sorbitol (Fig. 2C). We concluded that *pkd2* is an essential gene, required for cell growth and cell division.

Next, we attempted to construct a viable *pkd2* mutant. We replaced the endogenous *pkd2* promoter with a series of inducible promoters (Basi *et al.*, 1993; Maundrell, 1990). Among the three resulting *pkd2* mutants, the one with the strongest growth defect is *pkd2::P81nmt1-pkd2* (refer to as *pkd2-81KD*) (Fig. S2A), in which a weak inducible *81nmt1* promoter replaced the endogenous promoter. The *pkd2-81KD* mutant was also hypersensitive to high concentration of salts including CaCl_2_ (Fig. S2B). To quantify the depletion or over-expression of Pkd2p in the mutants, Pkd2p was tagged with GFP in both *pkd2-81KD* and *pkd2::P3nmt1-pkd2* to measure average numbers of Pkd2p molecules per cell (Fig. S2C and Movie S2). The number of Pkd2p molecules decreased by ∼70% in the *pkd2-81KD* cells but doubled in *pkd2::P3nmt1-pkd2*, when the *nmt1* promoters were suppressed (Fig. 2E). The depletion of Pkd2p led to strong morphological defects of *pkd2-81KD* cells, which could be reversed by removing thiamine to restore the expression of *pkd2* (Fig. 2F). In contrast, neither *trp663Δ* nor *trp1322Δ* cells exhibited an apparent morphological defect (Fig. S1B). We concluded that *pkd2-81KD* is a novel hypomorphic *pkd2* mutant that can be used in further studies of this gene.

We examined the *pkd2-81KD* mutant for its other defects. The mutant cells were significantly wider than the wild-type cells (Fig. S2D). The cell wall of this *pkd2* mutant, examined with transmission electron microscopy, was similar in appearance and thickness to that of the wild-type (Fig. 2G-H). Compared to the wild type cells, the actin cytoskeletal structures of the mutant appeared to be similar, but distribution of the interphase actin patches was less polarized in the mutant cells, suggesting a potential cell polarity defect (Fig. 2I). The requirement of Pkd2p was not limited to vegetative cells. The *pkd2* mutant spores initiated their outgrowth more slowly than the wild-type ones did (Fig. 2J), reminiscent of the defect observed in *pkd2Δ* spores noted above. This germination defect of the *pkd2-81KD* spores, was also apparent in their usual slow growth following tetrad dissections (data not shown). We concluded that Pkd2p likely plays an essential role in cell morphogenesis.

### Temporary deflation of the *pkd2* mutant cells

When examining *pkd2-81KD* cells with bright-field microscopy, we noticed that some of them appeared to be lysed. Since cell lysis is often associated with a defect in the cell wall biosynthesis, we characterized this phenotype in further details (Cortes *et al.*, 2012; Davidson *et al.*, 2016; Munoz *et al.*, 2013; Sethi *et al.*, 2016). To our surprise, these mutant cells didn’t lyse completely but they merely shrank temporarily, which we termed as “Deflation”, before quickly recovered. In all, 16% of the mutant cells underwent deflation and re-inflation (Fig. 3A and Movie S3). In contrast, no wild-type cell shrank under the same condition. The temporary deflation of the *pkd2* mutant cells lasted ∼20 mins (20 ± 8 mins, average ± S.D., n = 12) during which the cell size was reduced by ∼20% (Fig. 3D). Interestingly, the majority of deflated cells were not undergoing either cell division or septation (80%, n = 15), suggesting that neither triggered the deflation. To our knowledge, this unique phenotype has not been described before for any other mutant. The quick loss of cell volume and the following recovery to some extent resembled the response of wild-type cells to hypertonic stress.

**Figure 3:**
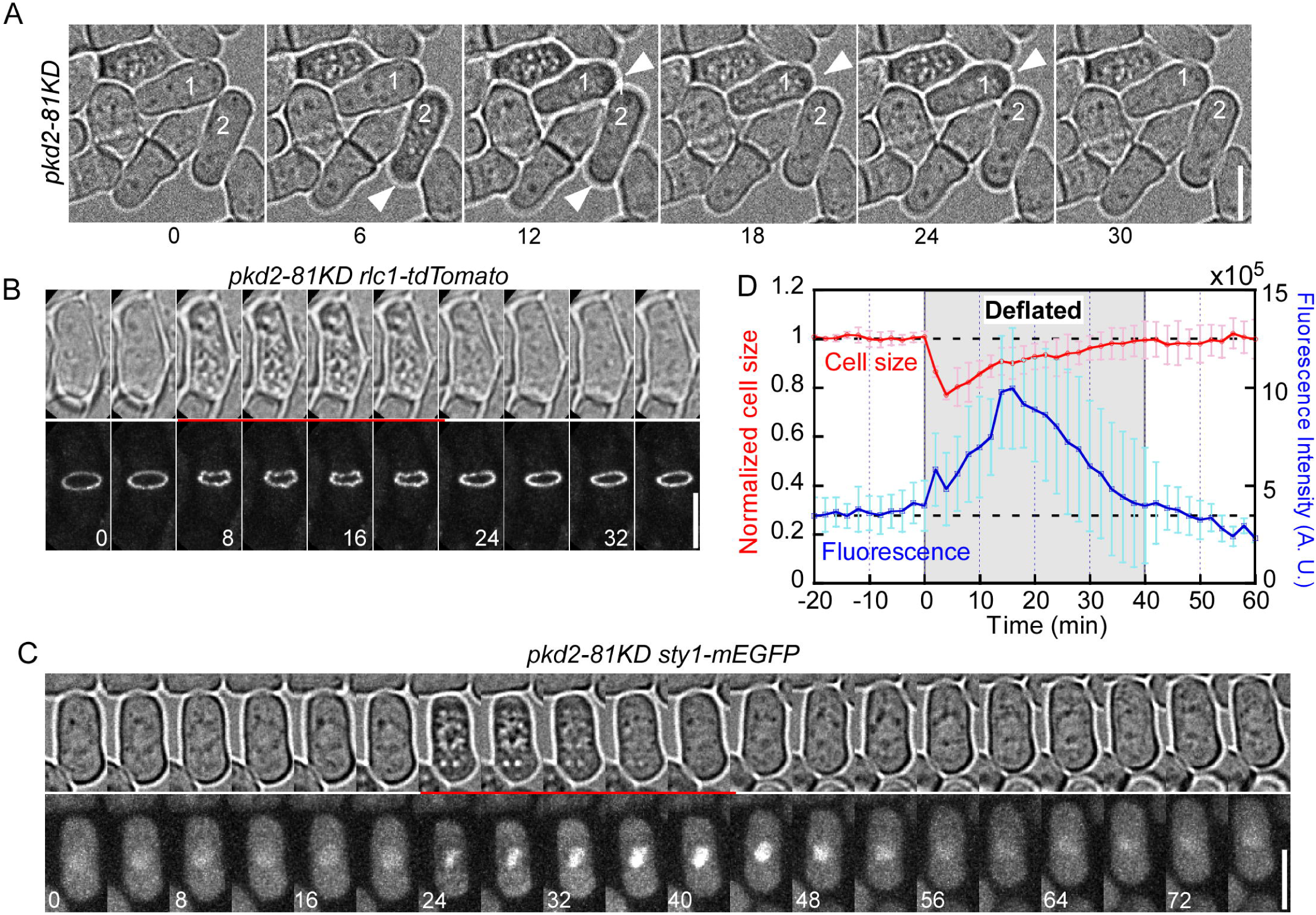
Deflation of the *pkd2* mutant cells. (A) Time-lapse micrographs of *pkd2-81KD* cells. Both cell 1 and 2 shrank temporarily (arrowheads). Numbers represent time in mins. (B) Time-lapse micrographs of a *pkd2-81KD* cell expressing Rlc1p-Tdtomato. The ring transformed into a “wavy band” (bottom) during the period of temporary deflation (top, red underline). (C) Time-lapse micrographs of a *pkd2-81KD* cell expressing Sty1p-GFP. Temporary shrinking (top, red underlined) of the cell was concomitant with increased Sty1p-GFP localization in the nucleus (bottom). Number represents time in mins. (D) Plot showing the time courses of average sizes (red line, normalized) and nuclear Sty1p-GFP fluorescence (blue line) of *pkd2-81KD* cells (n = 7). The shrinking reduced the average cell size by ∼20%. Concurrently, Sty1p-GFP fluorescence in the nucleus increased by about three-fold. Error bars represent standard deviations.

We determined whether deflation of the *pkd2* mutant cells was a result of osmotic stress by first examining turnovers of their cytoskeletal structures which would stop temporarily in response to osmotic stresses (Robertson and Hagan, 2008), We first examined the contractile ring and the actin patches, two actin cytoskeletal structures. Upon deflation of the mutant cells, their contractile ring stopped constricting abruptly and converted temporarily from a circular ring into a “wavy band” (Fig. 3B and Movie S4). The deflation triggered a stop in turnover of the endocytic actin patches as well (Fig. S3A). Not surprisingly, turnover of the microtubules stopped temporarily in the deflated cells too (Fig. S3B, Movie S5 and Movie S6). We concluded that turnover of the cytoskeletal structures temporarily was stopped temporarily in the deflated *pkd2* mutant cells as in osmotic stressed cells.

We next determined whether deflation of the *pkd2* mutant cells also activated the Sty1p-mediated MAPK pathway another hallmark of the cells under osmotic stresses. Activation of this MAPK pathway shuttles Sty1p into nucleus where the kinase mediates the stress-activated gene expressions (Degols *et al.*, 1996; Millar *et al.*, 1995; Shiozaki and Russell, 1995). We found significantly more Sty1p-GFP molecules in the nuclei of *pkd2-81KD* cells than the wild-type cells (Fig. S3C and S3D). When the *pkd2* mutant cells shrank, nuclear localization of Sty1p-GFP increased concomitantly by up to three folds, compared to pre-deflation (Fig. 3C, 3D and Movie S7). We concluded that the Sty1p-mediated MAPK pathway was hyper-activated during deflation of the *pkd2* mutant cells as was found in osmotically stressed cells. In addition, we also found that the deflation of *pkd2* mutant cells could be rescued by the osmotic stabilizer sorbitol (n > 500 cells) (Fig. S3E). In summary, we concluded that Pkd2p most likely regulates osmotic homeostasis during cell growth.

### Pkd2p regulates the ring closure during cytokinesis

To examine the role of Pkd2p during cytokinesis, we first measured the contractile ring assembly in *pkd2-81KD* cells. Overall, the ring assembled normally in the mutant cells (Movie S8 and S9). The *pkd2* mutation did not disrupt the placement of rings at the cell division plane (Fig. 4A). The rings were positioned properly in the *pkd2* mutant cells as well, unlike the oblique angled rings found in those mutants with a defect in the septum biosynthesis (Munoz *et al.*, 2013). We did notice that the mutant cells were significantly shorter than the wild-type cells during cytokinesis (Fig. 4B). The *pkd2* mutant cells also experienced a slight but significant delay in ring assembly and maturation, compared to the wild-type cells (29 ± 4 vs. 25 ± 4 mins, average ± S.D., P < 0.001) (Fig. 4C). We concluded that Pkd2p is not essential for either the assembly or maturation of the contractile ring.

**Figure 4:**
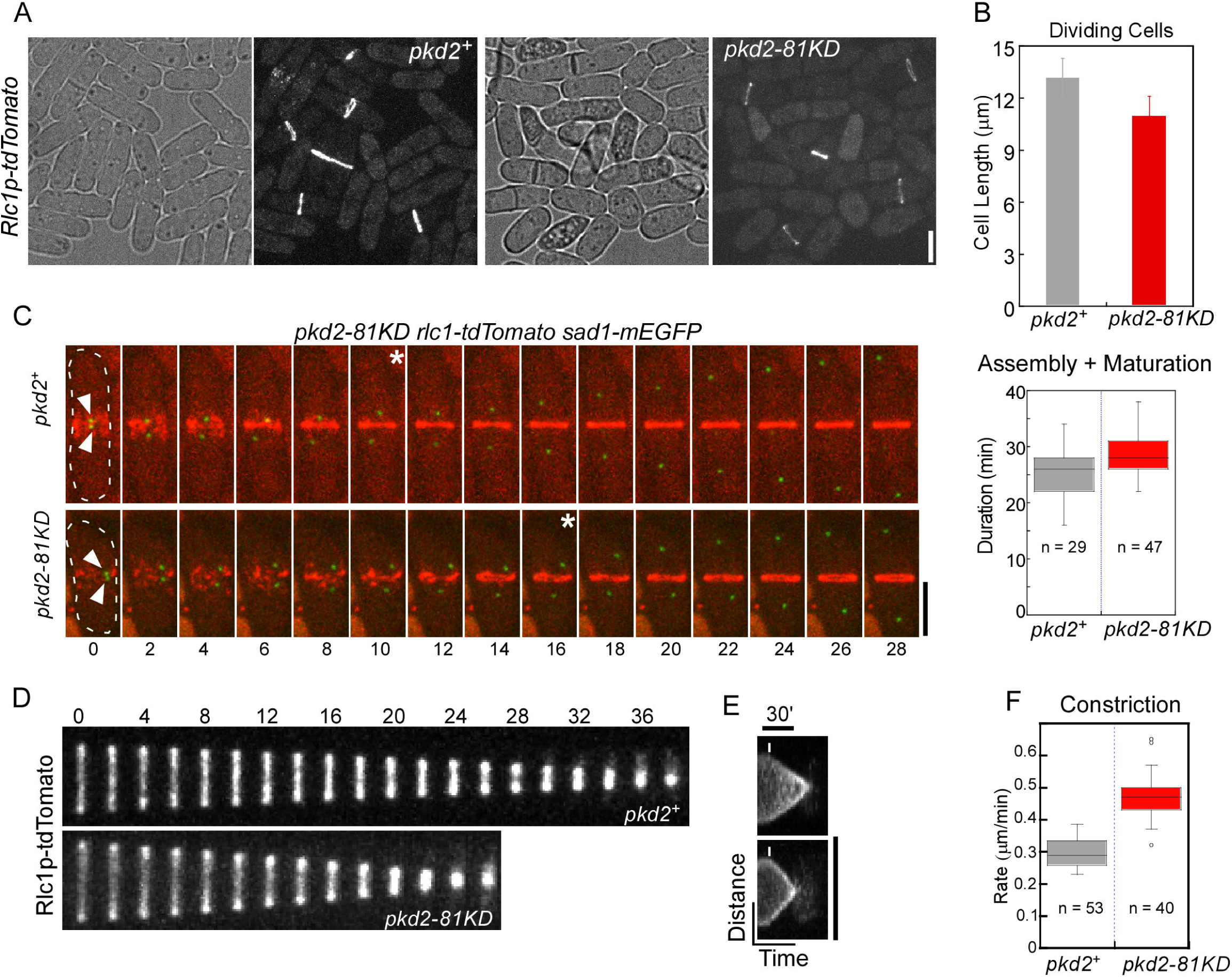
Increased ring closure rate in *pkd2-81KD* cells. (A) Micrographs of wild-type (*pkd2*^*+*^, left) and *pkd2-81KD* (right) cells expressing Rlc1p-tdTomato. Placement of the contractile rings in the mutant were similar to that in the wild-type cells. (B) Bar graph showing the average length of wild-type and *pkd2-81KD* cells during cytokinesis. The analysis included only cells with one complete contractile ring. The *pkd2* mutant cells were significantly shorter than the wild-type cells (n > 50, P < 0.001). (C) Ring assembly in *pkd2-81KD* cells. Left: Time-lapse fluorescence micrographs of a wild-type (top, *pkd2*^*+*^) and a *pkd2-81KD* cell (bottom). Both cells expressed Rlc1p-tdTomato (Red) and Sad1p-GFP (Green), a marker for SPBs. Ring assembly was slower in the mutant (∼6 mins) than the wild-type cells. Asterisks indicate the appearance of a complete ring. Numbers represent time in mins. Right: box plot showing the duration of ring assembly plus maturation in wild-type (grey) and *pkd2-81KD* cells (red). Ring assembly and maturation was slower in the mutant compared to the wild-type cells (P < 0.0001). (D-F) Ring closure in *pkd2-81KD* cells. Time-lapse fluorescence micrographs of the cell division plane in a wild-type (top) and a *pkd2-81KD* (bottom) cells. Frame interval is 2 min. The rings constricted more rapidly in the mutant than in the wild-type cells. (E) Fluorescence kymographs of ring closure in a wild-type (top) and a *pkd2* mutant (Bottom) cells. Lines: Duration of ring closure. (F) Box plot showing the average ring closure rates in wild-type (grey) and *pkd2-81KD* (red) cells. The rate increased by ∼50% in the mutant. Bars represent 5 µm. Error bars represent standard deviations.

Next, we determined whether Pkd2p regulates ring closure, a more likely role considering its localization during cytokinesis. Surprisingly, ring closure on average took less time in the mutant than in the wild-type cells (Fig. 4D), in spite of our earlier observation that the mutant cells were wider than the wild-type ones (Fig. S2D). As expected, the rings in the mutant cells contracted more than 50% faster than they did in the wild-type cells (0.47 ± 0.07 vs. 0.30 ± 0.04 µm/min, P < 0.0001) (Fig 4E and 4F) (Fig. S4A). Our measurement of ring closure in the wild-type cells, closely matched the previously published values (Chen and Pollard, 2011; Laplante *et al.*, 2015; Proctor *et al.*, 2012). We did not observe any sliding of the rings during their contraction in the mutant (n > 50), which would have occurred if the ring had not been anchored properly (Arasada and Pollard, 2014; Cortes *et al.*, 2012). Therefore, it was unlikely that this rapid ring closure in the mutant was a result of a failure to couple the ring to the plasma membrane or the cell wall. Unexpectedly, over-expression of Pkd2p did not slow the ring closure as one would had predicted (Fig. S4B and S4C), indicating that Pkd2p is necessary, but not sufficient in modulating the rate of ring closure. We concluded that Pkd2p modulates ring closure during cytokinesis but is dispensable for ring assembly, maturation or anchoring.

### Pkd2p is required for separation of daughter cells

We next examined the role of Pkd2p in cell separation, the final phase of fission yeast cytokinesis. Two lines of evidence indicated that Pkd2p may play an important role in cell separation. First, the fraction of *pkd2-81KD* cells with septa was about three times higher than the wild-type cells (Fig. 5A and 5B). In addition, close to 9% of the mutant cells were multi-septated (Fig. 5B). We also noticed that the septum in these multi-septated mutant cells frequently swung from one side to another at a rate of 0.05 ± 0.03 µm/min (n = 12). Secondly, the septum in many mutant cells were curved, deposited ectopically at the cell tips or unusually thick (Fig. 5A). We postulated that Pkd2p likely is essential for cell separation.

We examined separation of the *pkd2* mutant cells during cytokinesis with live microscopy. The wild-type cells reliably separated two daughter cells in ∼30 min (28 ± 5 mins, average ± S.D., n = 20) after completion of the ring closure (Fig. 5C). In contrast, the mutant cells took more than twice as long (75 ± 21 mins, n = 17) (Fig. 5D) with some of them taking as long as 120 min to separate (Fig. 5C). Furthermore, some mutant cells failed to separate completely, explaining the many multi-septated *pkd2* mutant cells. Wild-type cells typically separate symmetrically by detaching both sides of the new ends from each other simultaneously (Fig. 5C). In contrast, most mutant cells separated asymmetrically by usually detaching one side of the new ends first (Fig. 5C bottom row). We concluded that Pkd2p plays an essential role in separating the new ends during cell separation.

We next determined whether Pkd2p was required for either the septum biosynthesis or the cell wall degradation, two critical steps of cell separation. We first examined the localization of two glucan synthases Bgs1p and Bgs4p, both of which are required for the septum biosynthesis (Cortes *et al.*, 2007; Le Goff *et al.*, 1999; Liu *et al.*, 1999). Both GFP-Bgs1p and GFP-Bgs4p still localized to the cell division plane in the *pkd2-81KD* cells, as they did in the wild-type cells (Fig. 6A and 6B). Furthermore, the time course of recruitment and dispersal of either Bgs1p or Bgs4p at the division plane in the mutant remained largely unchanged as well (Fig. 6C and 6D). The septum did close more rapidly in the mutant compared to the wild-type cells, consistent with the quicker ring closure in the mutant (Fig. S4D and S4E). As a result, the number of Bgs1p molecules at the cell division plane of the mutant decreased slightly compared to the wild-type cells (Fig. 6C). In contrast, the number of GFP-Bgs4p molecules at the division plane rose slightly (Fig. 6D). We concluded that Pkd2p is dispensable for the localization of these two glucan synthases during cell separation. Next, we determined whether Pkd2p is required for the localization of Eng1p, a glucanase that degrades the septum at the end of cell separation (Martin-Cuadrado *et al.*, 2003; Sipiczki, 2007). The *pkd2* mutation did not alter the localization of Eng1p-GFP at the cell division plane (Fig. S4F and S4G). The number of Englp-GFP molecules at the division plane remained unchanged, compared to that in the wild-type cells. We concluded that Pkd2p is not required for the localization of Eng1p either and the mostly likely role of Pkd2p during cell separation is to regulate the turgor pressure in the separation of new ends.

**Figure 6:**
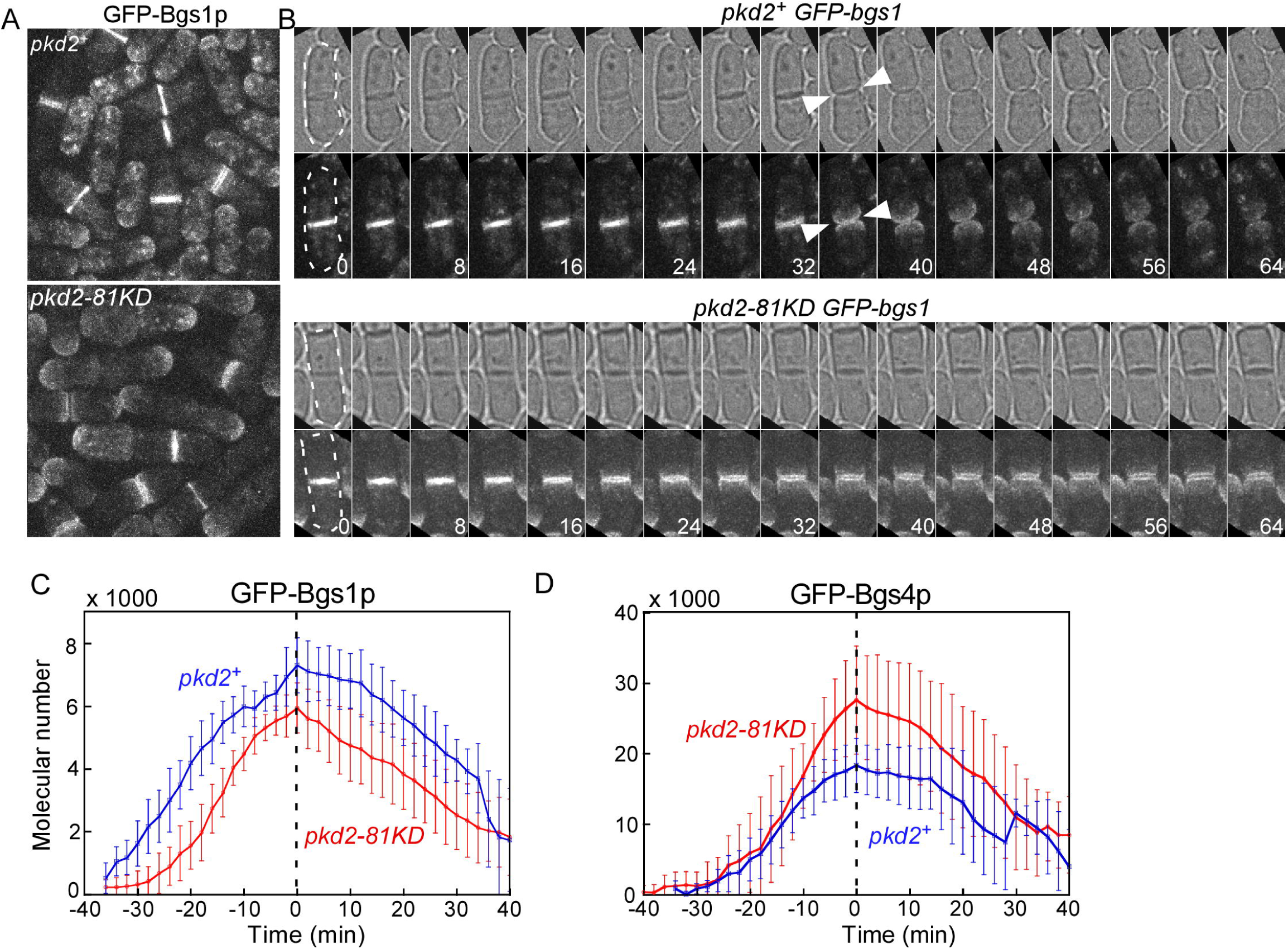
Localization of the glucan synthases Bgs1p and Bgs4p in *pkd2-18KD* cells. (A) Fluorescence micrographs of wild-type (*pkd2*^*+*^, top) and *pkd2-81KD* (bottom) cells expressing GFP-Bgs1p. The *pkd*2 mutation did not alter GFP-Bgs1p localization at the cell division plane. (B) Time-lapse micrographs of a wild-type (top) and a *pkd2-81KD* (bottom) cells expressing GFP-Bgs1p. The wild-type cells separated at 36 min (white arrowheads) after the ring closure (time zero), but the mutant cells failed to separate. Nevertheless, GFP-Bgs1p localization at the cell division plane appeared unchanged in the mutant. Frame interval is 4 min. (C-D) Line plots showing the time courses of GFP-Bgs1p (C) and GFP-Bgs4p (D) molecules localized at the division plane of either wild-type (blue line) or *pkd2-81KD* cells (red line). The number of GFP-Bgs1p molecules in the *pkd2* mutant was slightly reduced but the number of GFP-Bgs4p molecules was increased, compared to the wild-type cells (n > 5). Error bars represent standard deviations.

### Genetic interactions between *pkd-81KD* and the other cytokinesis mutants

Pkd2p localization at the cleavage furrow and its role in cell separation prompted us to examine whether it interacts with the other cytokinesis genes. We carried out a targeted screen to identify genetic interactions between *pkd2-81KD* and eleven other cytokinesis mutants (Fig. 7A). Four of them interacted negatively with the *pkd2* mutant (Fig. 7A), including the temperature sensitive mutants of *myo2* (type II myosin), *cdc12* (formin), *rng2* (IQGAP1 homologue) and *cdc15* (a F-BAR protein) respectively. All four genes are required for the ring assembly (Chang *et al.*, 1997; Eng *et al.*, 1998; Fankhauser *et al.*, 1995; Kitayama *et al.*, 1997). On the other hand, *cps1-191*, the temperature sensitive mutant of *bgs1*, exhibited no genetic interaction with *pkd2-81KD* mutant. We concluded that Pkd2p plays a synergistic role with Myo2p, Cdc12p, Rng2p and Cdc15p during cytokinesis.

**Figure 7:**
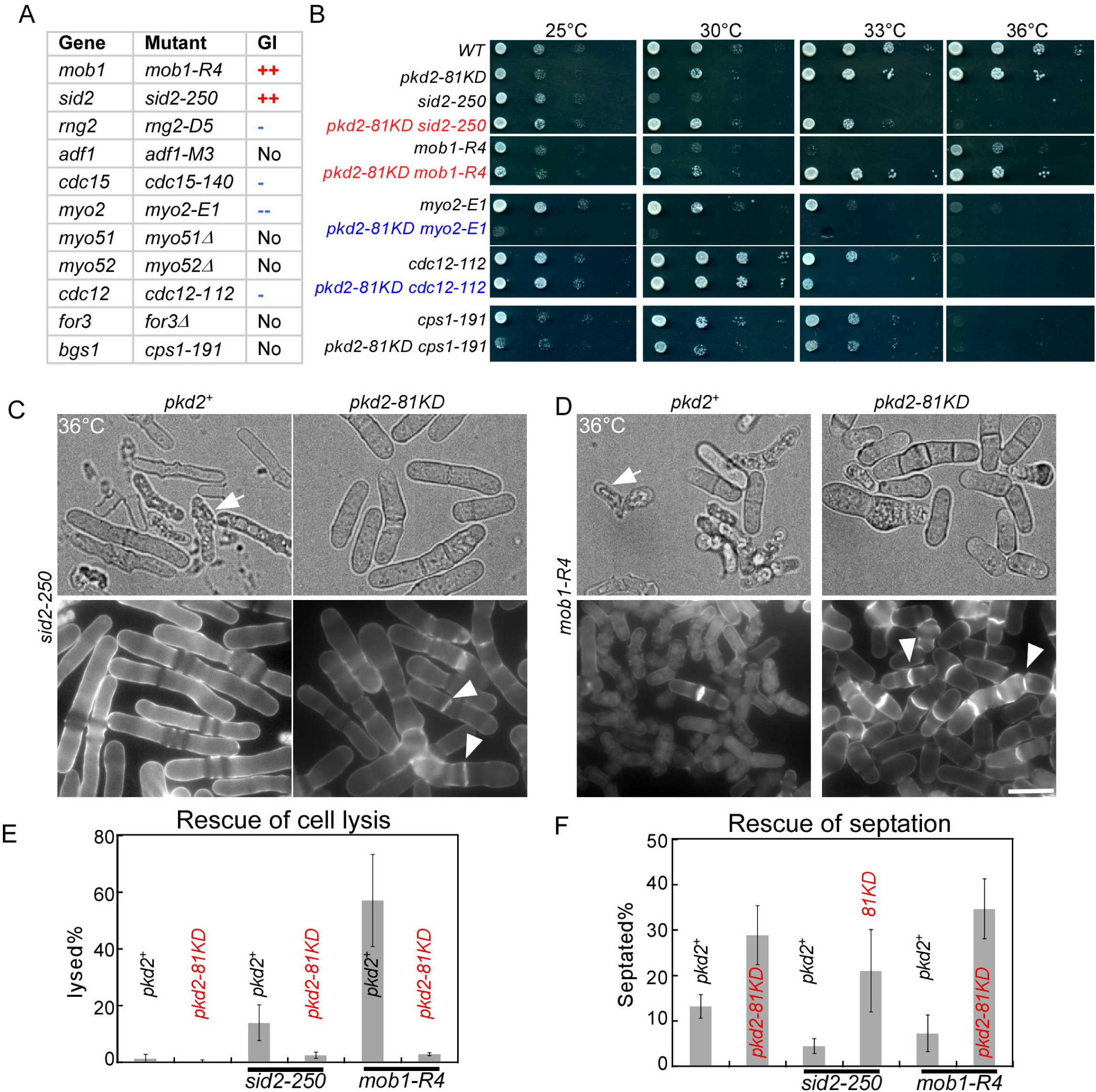
Genetic interactions between *pkd2-81KD* and the other cytokinesis mutants. (A) A table summarizing the genetic interactions between *pkd2-81KD* and eleven other cytokinesis mutants. Positive interactions are designated by either “++” (strong) while negative interactions are designated by either “--” (strong) or “-” (weak). “NO” indicates no genetic interactions. (B) Ten-fold dilution series of yeast cells at the indicated temperatures. *pkd2-81KD* mutant had positive genetic interactions with *mob1-R4* and *sid2-250* but negative interactions with *myo2-E1* and *cdc12-112*. It had no genetic interactions with *cps1-191*. (C-D) Micrographs of *sid2-250, sid2-250 pkd2-81KD, mob1-R4* and *mob1-R4 pkd2-81KD* cells grown. Top: Bright-field micrographs of the live cells. Bottom: fluorescence micrographs of the calcofluor-stained cells. Arrows: lysed cells. Arrowheads: the septum. The SIN mutants frequently lysed and few of them contained septum at the restrictive temperature, defects that were partially rescued by the *pkd2* mutation. (E-F) Bar graphs showing the percentage of either lysed (E) or septated cells (F) at 36°C. The *pkd2* mutation largely rescued both the lysis and septation defects of the SIN mutants. Error bars represent standard deviations.

Two SIN mutants, *mob1-R4* and *sid2-250*, had the strongest genetic interactions with *pkd2-81KD*. They are the temperature sensitive mutants of Sid2p kinase and its activator Mob1p respectively (Balasubramanian *et al.*, 1998; Hou *et al.*, 2000). Growth of *mob1-R4* was rescued by the *pkd2* mutation at the restrictive temperature (Fig. 7B). In comparison, the *pkd2* mutation only rescued the growth of *sid2-250* at semi-restrictive temperature. The SIN temperature-sensitive mutants failed in septation and eventually lysed, at the restrictive temperature (Balasubramanian *et al.*, 1998; Krapp *et al.*, 2004; Salimova *et al.*, 2000) (Fig. 7C and 7D). The *pkd2-81KD* mutation prevented the lysis of both SIN mutant cells (Fig. 7E and S5C). Additionally, the relatively mild septation defect of *mob1-R4* cells was also rescued by *pkd2-81KD* (Fig. 7F and S5D). In comparison, the septation defect of *sid2-250* mutant was only partially rescued by the *pkd2* mutation at both semi-restrictive and restrictive temperature (Fig. 7F and S5D). Additionally, the septum of *sid2-250 pkd2-81KD* cells were much thinner than those of wild-type cells (Fig. 7C). We concluded that Pkd2p and the SIN pathway play antagonistic roles in septation and maintaining cell integrity.

## Discussion

Forces, generated by the contractile ring, the septum and the turgor, are essential for fission yeast cytokinesis. However, we know relatively little about how these forces can be sensed and modulated by cells. Here we showed that a putative force-gated channel Pkd2p has several important functions during cytokinesis, including modulating the ring closure and promoting cell separation. Defects of the *pkd2-81KD* mutant cells suggest that the most likely role of this TRP channel is to regulate osmotic homeostasis. Not surprisingly, Pkd2p is well placed at the cleavage furrow during cytokinesis to execute this function. The genetic interactions between *pkd2-81KD* and the other cytokinesis mutants also support that this transmembrane protein may be a novel component of the cytokinetic machinery.

### Localization of Pkd2p channel at the cleavage furrow

To our knowledge, Pkd2p is the first putative ion channel that localizes to the cell division plane of fission yeast cells. Our discovery is compatible with the previous finding that Pkd2p was mostly found at the intracellular organelles and the plasma membrane (Aydar and Palmer, 2009; Palmer *et al.*, 2005). Pkd2p is one of the few TRP channels that have been found at the cleavage furrow so far. TRP3 channel of the zebrafish *Danio rerio*, which mediates the influx of calcium at the furrow, is the other notable example (Chan *et al.*, 2015). Many molecules in the contractile ring may interact with Pkd2p directly to recruit it to the cleavage furrow. This mechanism will be consistent with our observation that the contractile ring is required for Pkd2p localization. The septum may also interact with Pkd2p directly, considering that the initial appearance of Pkd2p at the furrow coincides with the start of primary septum synthesis. Future studies will be needed to determine the molecular mechanism of Pkd2p recruitment to the cell division plane.

Based on our study of fission yeast Pkd2p, it will be interesting to determine the localization of other Pkd2p homologues during cytokinesis. Budding yeast homologues of Pkd2p localize to both the ER and plasma membrane (Protchenko *et al.*, 2006) but their localization during cytokinesis has not been examined. Among the two human Pkd2p homologues, PKD1 is primarily localized at the plasma membrane (Hughes *et al.*, 1995) and PKD2 is mostly found on the ER membrane (Gonzalez-Perrett *et al.*, 2001), although PKD2 does localizes to the mitotic spindles through its interaction with formin mDia1 in cultured kidney epithelial cells (Rundle *et al.*, 2004). Additionally, most Pkpd2 homologues, including those of humans, *C. elegans* and *D. melanogaster*, also localize to the cilia of non-dividing cells (Barr *et al.*, 2001; Barr and Sternberg, 1999; Gao *et al.*, 2003; Nauli *et al.*, 2003; Pazour *et al.*, 2002; Watnick *et al.*, 2003; Yoder *et al.*, 2002). Future work would be necessary to determine the cytokinetic localization of Pkd2p in dividing animal cells.

### Molecular functions of Pkd2p during cytokinesis

Our study suggests that Pkd2p most likely regulates osmotic homeostasis during cytokinesis as well as cell morphogenesis. This model is supported by at least three lines of evidence. First, the unique deflation defect of the *pkd2* mutant could be due to a defect in the regulation of turgor pressure. The temporary shrinking of *pkd2-81KD* cells activated the stress response pathway mediated by Sty1p. The shrinking also stopped turnovers of all cytoskeletal structures that we examined. All of these phenotypes are consistent with mis-regulated turgor pressure. In addition, the septa of multi-septated *pkd2-81KD* cells often swung from side to side, suggesting that turgor pressure fluctuates drastically in these mutant cells. Secondly, the morphological defects of *pkd2-81KD* cells also imply a defect in the osmotic regulation during cell morphogenesis. Compared to wild-type cells, the *pkd2* mutant cells are shorter but wider and the distribution of their actin patches is partially depolarized. Furthermore, both *pkd2Δ* and *pkd2-81KD* spores either failed or delayed their outgrowth during germination. These observations are consistent with a defect in the regulation of turgor pressure, that is essential to fission yeast cell growth (Atilgan *et al.*, 2015; Davi *et al.*, 2018). Third, the cytokinesis defects of *pkd2-81KD* cells are also consistent with mis-regulated turgor pressure. Both rapid ring and septum closure in the mutant cells can be a result of low turgor pressure which could reduce resistance to the furrow ingression and increase the ring closure rate, compared to the wild type cells. This hypothesis is consistent with the important role of turgor in determining ring closure rate (Proctor *et al.*, 2012). On the other hand, slow or failed separation of the *pkd2* mutant cells may be attributed to mis-regulated turgor at the cell division plane (Abenza *et al.*, 2015; Atilgan *et al.*, 2015; Sipiczki, 2007). Although future studies will be necessary to directly measure the putative abnormities of turgor pressure in the *pkd2* mutation cells, our observations strongly support that Pkd2p plays an essential role in osmotic homeostasis during cytokinesis and cell morphogenesis.

As a TRP channel, Pkd2p could potentially regulate the turgor pressure through a number of ways. It may control the ion influx to directly modulate the turgor. As a permissive cation channel, Pkd2p could allow the calcium influx to regulate the turgor pressure, considering its localization on the plasma membrane (Palmer *et al.*, 2005). In this way, Pkd2p may be similar to another TRP channel TRPV4 that regulates osmolarity of animal cells (Colbert *et al.*, 1997; Liedtke and Friedman, 2003; Strotmann *et al.*, 2000). Alternatively, Pkd2p may regulate the intracellular osmolarity indirectly through interacting with the other signaling pathways including the SIN pathway. It is worth noticing that both Pkd2p and the Sid2p/Mob1p complex localize to the cell division plane during late cytokinesis (Goss *et al.*, 2014; Sparks *et al.*, 1999). More studies will be needed to determine how Pkd2p acts through these potential mechanisms to regulate the turgor pressure.

### Molecular functions of the Pkd2p family proteins

Our study identified a novel function of the fungal Pkd2p family proteins in regulating cytokinesis. This family of transmembrane proteins can be found among many fungi species and is one of seventeen essential fungal protein families (Hsiang and Baillie, 2005). Four *Saccharomyces cerevisiae* Pkd2 homologues, Flc1 (<u>Fl</u>avin <u>c</u>arrier 1), Flc2, Flc3 and YOR365c, have been identified. Unlike fission yeast *pkd2*, none of these budding yeast homologues is essential (Protchenko *et al.*, 2006; Rigamonti *et al.*, 2015; Vazquez *et al.*, 2016). However, deletion of both *flc1* and *flc2* is lethal. These four proteins appear to play a role in osmotic regulation (Protchenko *et al.*, 2006), heme transportation (Protchenko *et al.*, 2006) and calcium homeostasis (Rigamonti *et al.*, 2015), although the molecular mechanism of their functions is unknown. It also remains to be determined whether any of these four budding yeast genes is required for cytokinesis. Other fungal Pkd2p homologues include s*pray* of *Neurospora crassa* (Bok *et al.*, 2001), FlcA/B/C of *Aspergillus fumigatus* (de Castro *et al.*, 2017), and CaFlc1-3 of *Candida albicans* (Protchenko *et al.*, 2006). Interestingly, all of them regulate hyphal growth of the fungi, similar to the fission yeast Pkd2p regulation of cell growth. Our proposed role of Pkd2p in regulating osmotic homeostasis would be consistent with the central role of turgor in the fungal growth(Lew, 2011).

In multicellular eukaryotic organisms, Pkd2p is required for several developmental processes but the underlying molecular mechanism is far from clear. Both *D. melanogaster* and *C. elegans* homologues of Pkd2p contribute to the locomotion of sperms and the *pkd2* mutations lead to male sterility (Barr and Sternberg, 1999; Gao *et al.*, 2003; Watnick *et al.*, 2003). The human Pkd2p homologues PKD1 and PKD2 are required for cardiac development and left-right orientation (Pennekamp *et al.*, 2002; Wu *et al.*, 2000). Their function in kidney has been studied most extensively, because loss of function mutations of either genes result in a hereditary human disease, ADPKD (for review see (Chapin and Caplan, 2010)). This renal disorder affects more than 600,000 Americans alone. It is characterized by progressive proliferation of liquid-filled cysts in the kidney that often culminates to a terminal renal failure. (Mochizuki *et al.*, 1996)Despite years of work, it still remains largely unknown how the mutations of these human Pkd2p homologues lead to ADPKD. Study of fission yeast Pkd2p and its function in osmotic homeostasis will likely provide us with critical insights into the molecular functions of human PKD1 and PKD2 as well as the pathogenesis of ADPKD.

## Materials and Methods

### Yeast genetics

Yeast cell culture and genetics procedures were carried out according to the standard methods. A SporePlay+ dissection microscope (Singer, England) was used for tetrad dissections. Most yeast strains were constructed through PCR based homologous recombination (Bahler *et al.*, 1998). To construct *trp663::trp663-GFP, trp1322::trp1322-GFP and pkd2::pkd2-GFP*, we integrated the mEGFP sequence at the endogenous locus of each gene. We constructed *trp1322*^*+*^/*trp1322Δ* by removing one copy of the ORF. The diploid cells were then sporulated and selected to isolate *trp1322Δ*. The *trp663Δ* mutant is from a fission yeast deletion library (Bioneer, South Korea) and was confirmed by PCR. To examine the viability of the *pkd2Δ* mutant, we constructed the diploid *pkd2*^*+*^*/pkd2Δ* by deleting one copy of the ORF through PCR-based homologous recombination. The diploid was then sporulated and dissected into more than 20 tetrads which were deposited onto either YE5s or YE5s plus 1.2M sorbitol agar plates. For microscopy of *pkd2Δ* spores, a small block of agar containing the spore was removed from the agar plate and deposited on a glass-bottomed petri dish for observation. To replace the endogenous *pkd2* promoter, we integrated inducible *nmt1* promoters at the endogenous locus, preceding the start codon of the *pkd2* ORF. To construct *pkd2::P81nmt1-pkd2-GFP-ura*, we constructed *pkd2::pkd2-GFP-ura* first, before replacing the endogenous promoter with *P81nmt1*. The resulting yeast strains were confirmed by both PCR and Sanger sequencing method. The *pkd2-81KD* spores were obtained by digesting the tetrads with Glusulase II (PerkinElmer, USA) and observed under a YE5s agar pad for 8-12 hrs with bright-field microscopy.

### Microscopy

For microscopy, exponentially growing yeast cells at 25°C with a density between 5.0*10^6^/ml to 1.0*10^7^/ml in YE5s liquid media (unless specified), were harvested by centrifugation at 4,000 rpm for 1 min and re-suspended in YE5s. The re-suspended cells were then applied to a 25% gelatin + YE5s pad, sealed under the coverslip with VALEP (a mix of an equal amount of Vaseline, lanolin and paraffin). Live microscopy was carried out on an Olympus IX71 microscope equipped with both 100 × (NA = 1.41) and 60 × (NA = 1.40) objective lens, a confocal spinning disk unit (CSU-X1, Yokogawa, Japan), a motorized XY stage and a Piezo Z Top plate (ASI, USA). The images were acquired on an Ixon-897 EMCCD camera controlled by iQ3.0 (Andor, Ireland). Solid-state lasers of 488 nm and 561 nm were used in the confocal fluorescence microscopy, at a power of no more than 5 mW (< 10%). Unless specified, we imaged cells by acquiring a Z-series of 15 slices at a step-size of 0.5 µm. Live microscopy was conducted in a room where the temperature was maintained at around 20°C. To minimize the temperature variations, we usually imaged both wild-type and the *pkd2* mutant cells on the same date. For temperature sensitive mutants, we inoculated the cultures at the restrictive temperatures for 4 hours before either imaging them at room temperature immediately or fixing the cells before imaging. For visualizing cell wall, we stained cells with 10 µg/ml of calcofluor (Sigma) and used an Olympus IX81 microscope equipped with a CCD camera and a mercury lamp for the epifluorescence microscopy. For visualizing the actin cytoskeletal structure, we fixed cells with 16% EM-grade paraformaldehyde (EMS, USA) and stained them with Bodipy-phallacidin (Invitrogen, USA), as described previously (Chen *et al.*, 2014).

### Electron microscopy

Transmission electron microscopy was carried out at the Microscopy and Image Analysis Laboratory core at the University of Michigan (Ann Arbor, MI). Briefly, exponentially growing cells were harvested, rinsed with 0.1M phosphate buffer (PH = 7.4) before being fixed with 2.5% glutaraldehyde (EMS, USA). The fixed cells were stained with 1% osmium tetroxide to increase contrast and embedded in epoxy resin overnight at 4°C. The samples were then sectioned at a thickness of 70 nm (Leica EM UC7). The EM specimens were imaged with a JEM-1400 (Joel, Japan) transmission electron microscope. The thickness of cell wall was measured as the average distance between the plasma membrane and outer boundary of the cell wall, at more than ten randomly selected sites.

### Image processing

We used Image J (NIH) to process all the images, with either freely available or customized macros/plug-ins. For quantitative analysis, the fluorescence micrographs were corrected for X-Y drifting using StackReg plug-in (Thevenaz *et al.*, 1998) and for photo-bleaching using EMBLTools plug-in (Rietdorf, EMBL Heidelberg). Average intensity projections of Z-slices were used for quantifications. The contractile ring localization of Pkd2p-GFP was quantified by measuring the GFP fluorescence in a 3.6 µm by 0.8 µm (36 by 8 pixels) rectangle centering on the cell division plane, corrected with background subtraction through measuring the background fluorescence intensities in two 3.6 µm by 0.2 µm rectangle (36 by 2 pixels) adjoined to the division plane. The cell width was measured using both live and fixed cells, which yielded similar results. The length of cytokinetic cells was measured using bright field images of the live cells expressing Rlc1p-tdTomato.

The nuclear localization of Sty1p-GFP was measured by quantifying GFP fluorescence intensities in the nuclei with background subtraction. To simplify the analysis, we assumed the nucleus as a circle of 2 µm diameter that centers on the nucleus localization of Sty1p-GFP. The background fluorescence was determined by measuring the fluorescence intensities of Sty1p-GFP in the cytoplasm surrounding the nucleus. Total area of a cell was measured based upon the cytoplasmic fluorescence of Styp1-GFP after the cell was segmented semi-automatically using Binary conversion and Analysis Particles tools of Image J.

The rate of ring closure is measured by analyzing fluorescence kymographs of a ring during cytokinesis. The kymograph was constructed from the time-lapse fluorescence micrographs (Fig. 4D), which was then used to determine both the start and end of ring closure. The diameter of a ring was determined at the start of ring closure. For quantitative fluorescence microscopy, our spinning-disk confocal microscope was calibrated using a previously published method (Wu and Pollard, 2005). Briefly, we imaged wild-type and seven fission yeast strains expressing GFP tagged proteins to produce a calibration curve (R^2^ > 0.95). Slope of the calibration curve was used to determine the ratio of fluorescence intensities to molecule numbers. The figures were made with Canvas X (ACDsee systems, Canada). The domain structure of Pkd2p is based on the prediction by Pfam and NCBI. The predicted topology of Pkd2p was created by Protter (http://wlab.ethz.ch/protter/), a protein topology data aggregation server (Omasits *et al.*, 2014).

## Acknowledgement

Q.C. designed the study. Z.M., D.S., A.P., B.M., and Q.C. carried out the experiments. Q.C. wrote the manuscript with inputs from Z.M., D.S. and A.P.. This work was supported by the University of Toledo Start-up Fund and deArce-Koch Memorial Endowment Fund to Q.C.. We would like to thank the groups of Carlos R. Vázquez de Aldana (IBFG, Spain), Juan Carlos Ribas (University of Salamanca, Spain), Dannel McCollum (University of Massachusetts Medical School) and Jianqiu Wu (Ohio State University) for generously sharing yeast strains with us. The authors expressed their appreciation of the help from their colleagues at the Department of Biological Sciences, Richard Komuniecki and Song-Tao Liu, in the preparation of this manuscript.

---

**Figure 5: Separation defect of *pkd2-81KD* cells during cytokinesis.** (A) Fluorescence micrographs of calcofluor-stained wild-type (*pkd2*^*+*^, Left) and *pkd2-81KD* (Right) cells. Septum of the mutant cells were often curved (arrow), thicker than normal (asterisk) or deposited ectopically (arrowhead). (B) Bar graph showing the percentage of wild-type and *pkd2-81KD* cells with at least one septa (n > 500). The percentage of multi-septated cells is shown in red. The fraction of septated *pkd2-81KD* cells was almost three times higher than that of the wild-type cells. (C-D) Separation defect of the *pkd2* mutant cells. (C) Time-lapse micrographs of a wild-type (Top) and two *pkd2-81KD* (Middle and Bottom) cells. All three cells expressed Rlc1p-tdTomato. Left: fluorescence micrographs of the cell when the ring closure was completed (time zero). Numbers represent time in mins. Frame intervals are 4 (top), 6 (middle) and 10 (bottom) mins respectively. The wild-type cell took 32 mins to separate (Red arrowheads). In contrast, the two *pkd2-81KD* cells took 60 and 120 mins respectively. (D) Dot plot showing the durations of cell separation in wild-type and *pkd2-81KD* cells (n > 15). The horizontal lines represent averages. The mutant cells separated more slowly than the wild-type cells.

**Figure S1.**
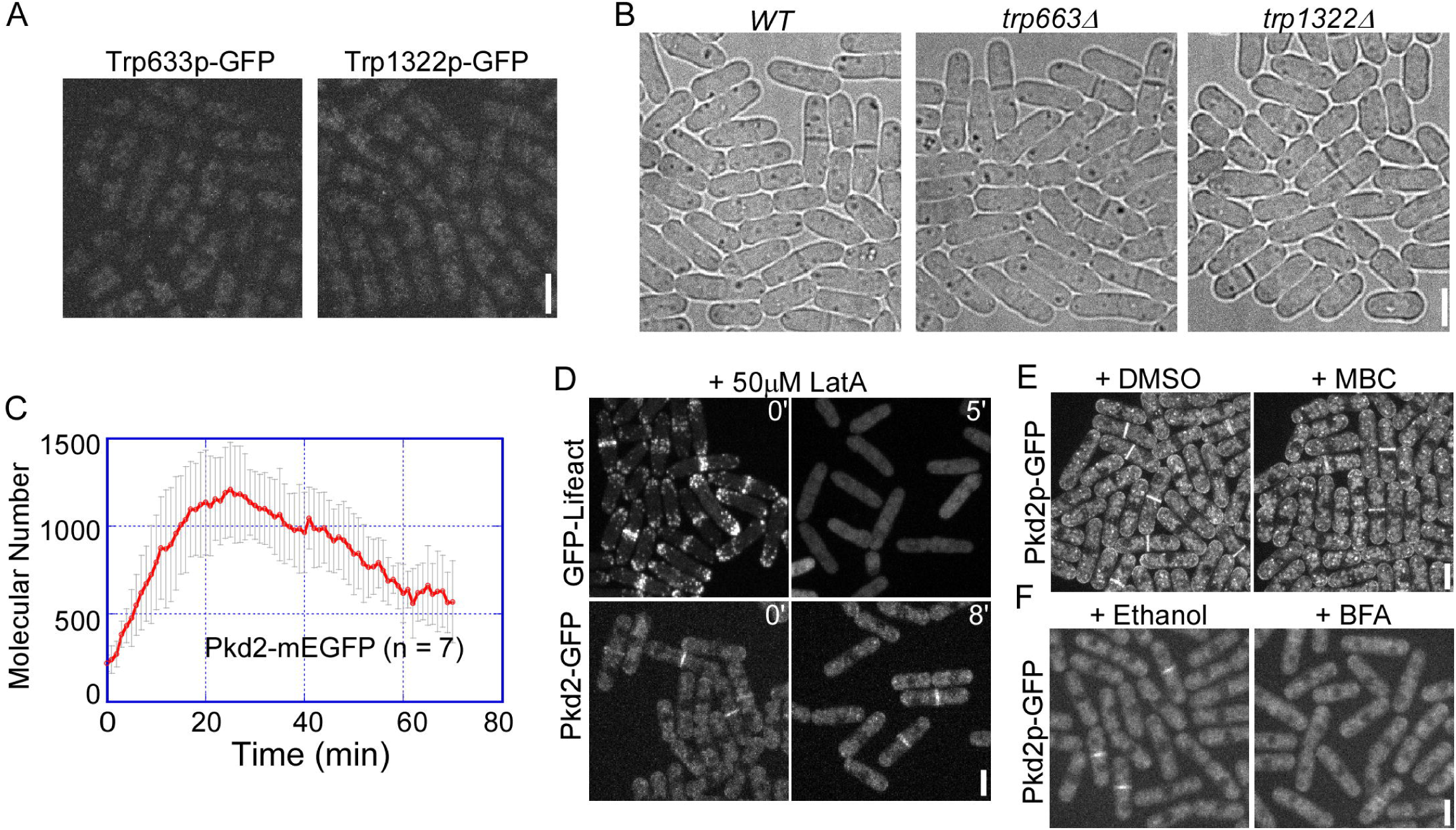
The localization of Pkd2p-GFP at the cell division plane, related to Fig. 1. (A) Fluorescence micrographs of cells expressing either Trp663p-GFP (left) or Trp1322p-GFP (right). Neither proteins exhibited significant localization at the cell division plane. (B) Bright-field micrographs of wild-type (*WT*), *trp663Δ* and *trp1322Δ* cells at 25°C. Neither mutants exhibited a significant morphological defect. (C) Line plot showing the time course of average numbers of Pkd2p-GFP molecules at the cell division plane. The number of Pkd2p-GFP molecules peaked at ∼ 1,200. (D) Fluorescence micrographs of the 50 µM LatA treated cells that expressed either GFP-Lifeact (top) or Pkd2p-GFP (bottom). The actin cytoskeletal structures were dissembled by LatA in less than 5 mins but Pkd2p-GFP remained at the division plane. (E) Fluorescence micrographs of cells treated for 60 mins with either DMSO (control) or 50 µM MBC, a microtubule-depolymerizing drug. Pkd2p-GFP maintained its localization at the division plane when the microtubules were depolymerized. (F) Fluorescence micrographs of cells treated for 70 min with either ethanol (control) or 100 µg/ml BFA, an inhibitor of the secretory pathway. Pkd2p-GFP was displaced from the division plane in the presence of BFA.

**Figure S2.**
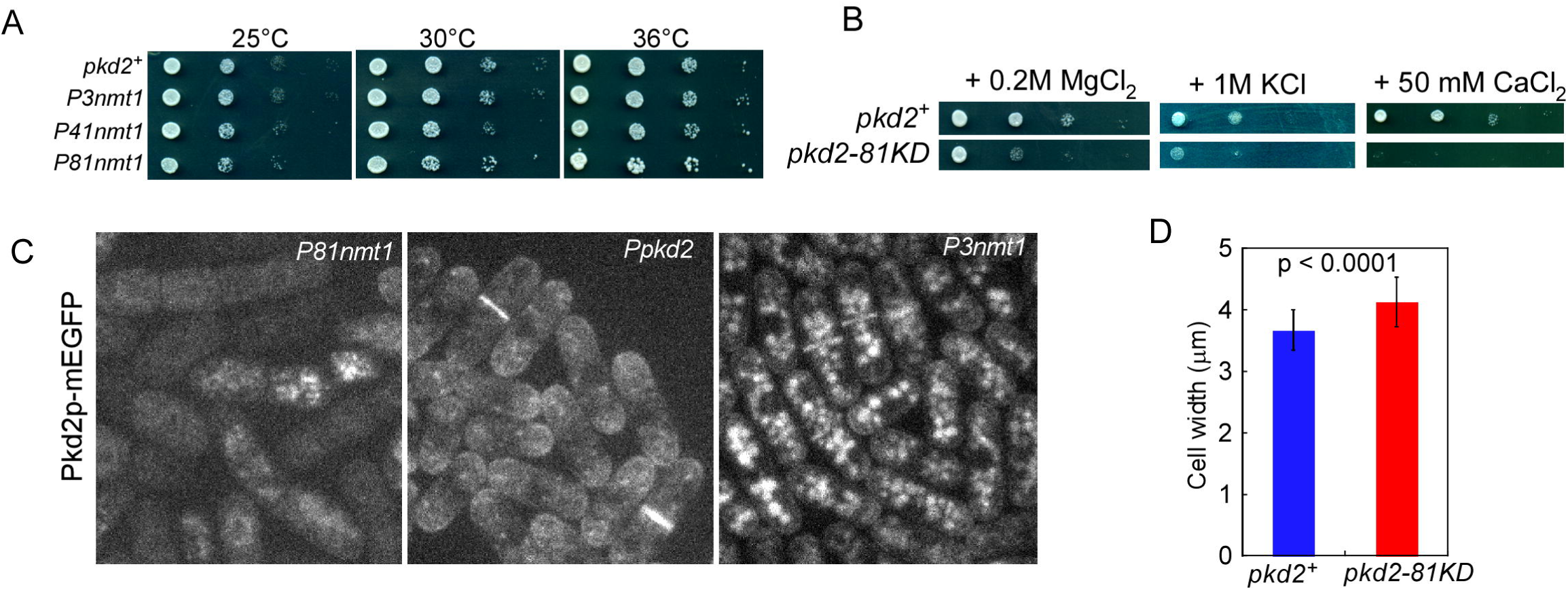
Pkd2p is an essential protein required for both cell growth and cell division, related to Fig. 2. (A) Ten-fold dilution series of wild-type and three *pkd2* mutants, *pkd2::P3nmt1-pkd2 (P3nmt1-pkd2), pkd2::P41nmt1-pkd2 (P41nmt1-pkd2) and pkd2::P81nmt1-pkd2 (P81nmt1-pkd2).* Among the three mutants, only *pkd2::P81nmt1-pkd2* exhibited a growth defect. (B) Ten-fold dilution series of wild-type and *pkd2-81KD* cells grown on YE5s plates plus the salts. The plates were incubated at 30°C for 2 days. The *pkd2* mutant was hypersensitive to MgCl_2_, KCl and CaCl_2_. (C) Fluorescence micrographs of *pkd2::P81nmt1-pkd2-GFP* (P81nmt1), *pkd2::Ppkd2-pkd2-GFP* (Ppkd2), and *pkd2:P3nmt1-pkd2-GFP* (P3nmt1) cells in YE5s. Average intensity projections are shown. Pkd2p-GFP fluorescence decreased in the *pkd2::P81nmt1-pkd2-GFP* cells but increased in the *pkd2::P3nmt1-pkd2-GFP* cells. (D) Bar graph showing the average width of wild-type (*pkd2*^*+*^) and *pkd2-81KD* cells (n > 50). The mutant cells were ∼10% wider. Error bars represent standard deviations.

**Figure S3.**
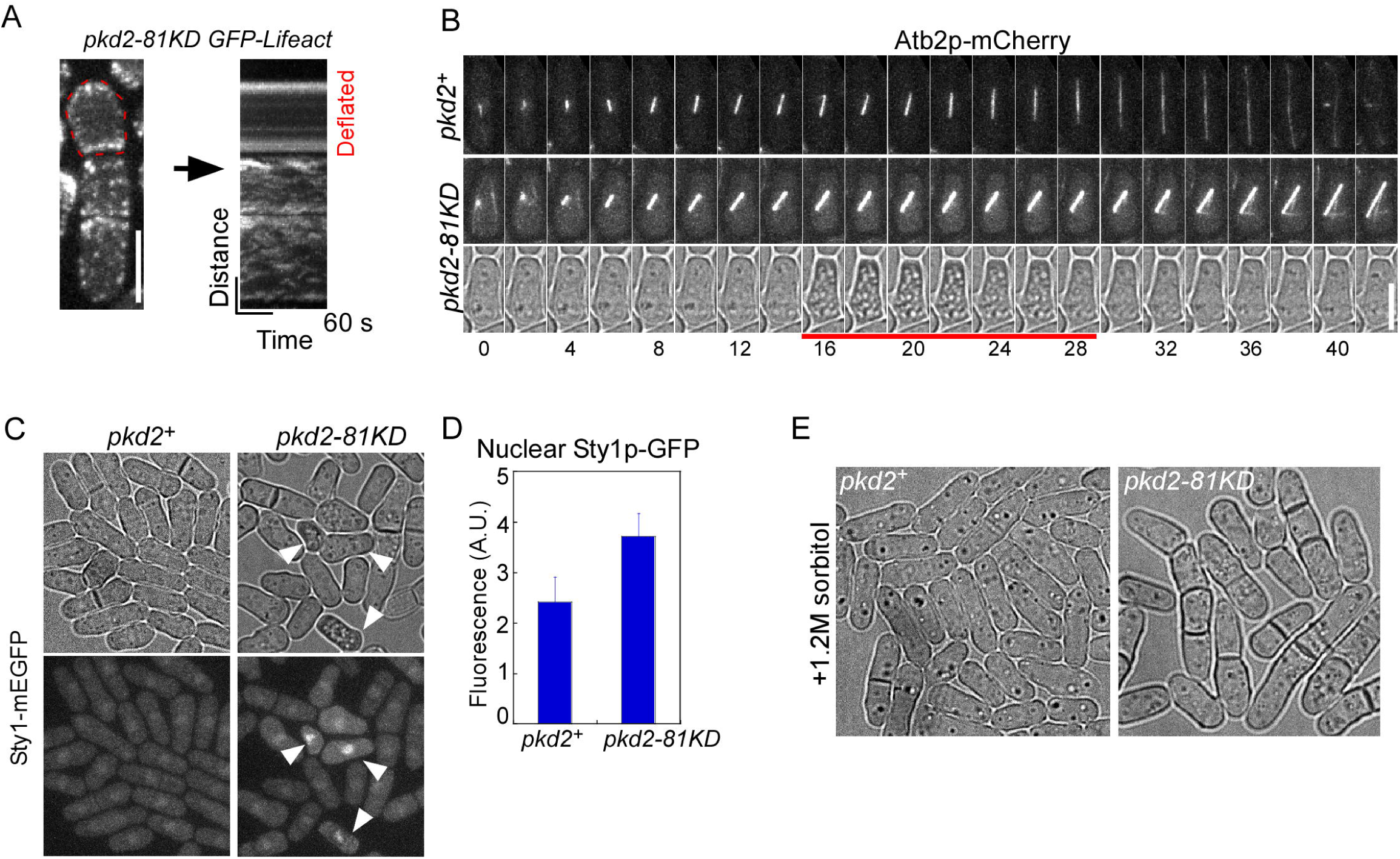
Temporary deflation of the *pkd2* mutant cells, related to Fig. 3. (A) Turnover of the actin patches in a deflated *pkd2-81KD* cell. Left: fluorescence micrograph of three GFP-Lifeact expressing *pkd2-81KD* cells separated by septum. The cell on the top shrank temporarily while the bottom two did not. Right: fluorescence kymograph of the three cells (60s). Turnover of the actin patches paused in the deflated cell (top, straight lines) but it continued in the other cells (bottom, discontinued stripes). (B) Time-lapse fluorescence micrographs of a wild-type (*pkd2*^*+*^, top) and a *pkd2-81KD* cells (middle and bottom). Both expressed Atb2p-mCherry as a marker for microtubules. Numbers represent time in mins. The mitotic spindles stopped elongating in the *pkd2-81KD* cell during the temporary deflation (bottom, red underline). (C) Micrographs of wild-type (*pkd2*^*+*^, left) and *pkd2-81KD* cells (right) expressing Sty1p-GFP. More Sty1p-GFP fluorescence was found in the nuclei of *pkd2-81KD* cells (arrowheads), compared to the wild-type cells. (D) Bar graph showing the average intensities of nuclear Sty1p-GFP fluorescence in wild-type and *pkd2-81KD* cells (n > 60). Sty1p-GFP fluorescence increased significantly in the nucleus of mutant cells (P < 0.001). Error bars represent standard deviations. (E) Micrographs of wild-type (*pkd2*^*+*^) and *pkd2-81KD* cells in YE5s plus 1.2 M sorbitol. The osmotic stabilizer rescued the deflation defect of mutant cells.

**Figure S4.**
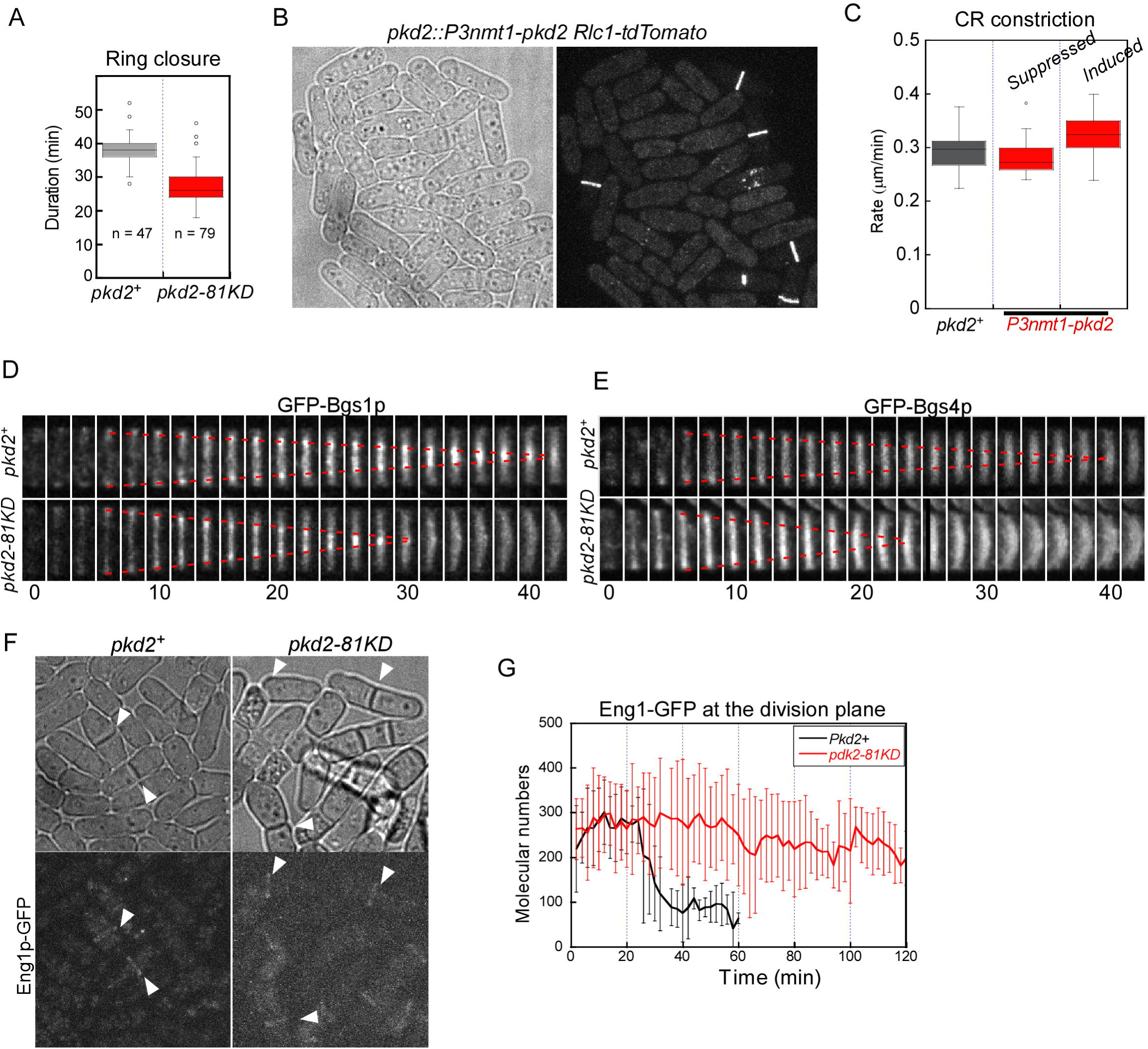
Cytokinesis defects of *pkd2-81KD* cells, related to Fig. 4, 5 and 6. (A) Box plot showing the duration of ring closure in wild-type (*pkd2*^*+*^) and *pkd2-81KD* cells. The ring closure took less time in the mutant cells. (B-C) Over-expression of Pkd2p did not lead to a cytokinesis defect. (B) Micrographs of *pkd2::P3nmt1-pkd2* cells expressing Rlc1p-tdTomato. Over-expression of Pkd2p did not alter either the placement or assembly of contractile rings. (C) Box plot showing the rates of ring closure in wild-type (*pkd2*^*+*^) and *pkd2::P3nmt1-pkd2* cells (n > 15). Over-expression of Pkd2p, either in EMM5s (inducing) or in YE5s (suppressing), did not alter the ring closure rate significantly, compared to the wild type (P > 0.05). (D-E) Time-lapse fluorescence micrographs of the septum in wild-type (*pkd2*^*+*^, top) and *pkd2-81KD* (bottom) cells. The cells expressed either GFP-Bgs1p (D) or GFP-Bgs4p (E). Red dash lines trace the septum closure over time. The localization of either Bgs1p or Bgs4p at the septum were not altered in the *pkd2* mutant, although the septum closure became more rapid in the mutant, compared to wild-type cells. Numbers represent time in mins. Frame interval is 2 min. (F-G) Pkd2p is not required for the localization of the glucanase Eng1p at the cell division plane. (F) Micrographs of wild-type (*pkd2*^*+*^, left) and *pkd2-81KD* (right) cells expressing Eng1p-GFP. The *pkd2* mutation did not alter Eng1p-GFP localization at the cell division plane (white arrowheads). (G) Line plot showing the time course of average numbers of Eng1p-GFP molecules at the division plane in wild-type (Black line) and *pkd2-81KD* (Red line) cells (n > 5). The *pkd2* mutation did not alter the molecular numbers of Eng1p-GFP at the division plane. Eng1-GFP remained at the division plane of the mutant cells after failed cell separation. In comparison, Eng1p diffused away from the division plane of the wild-type cells after the separation. Error bars represent standard deviations.

**Figure S5.**
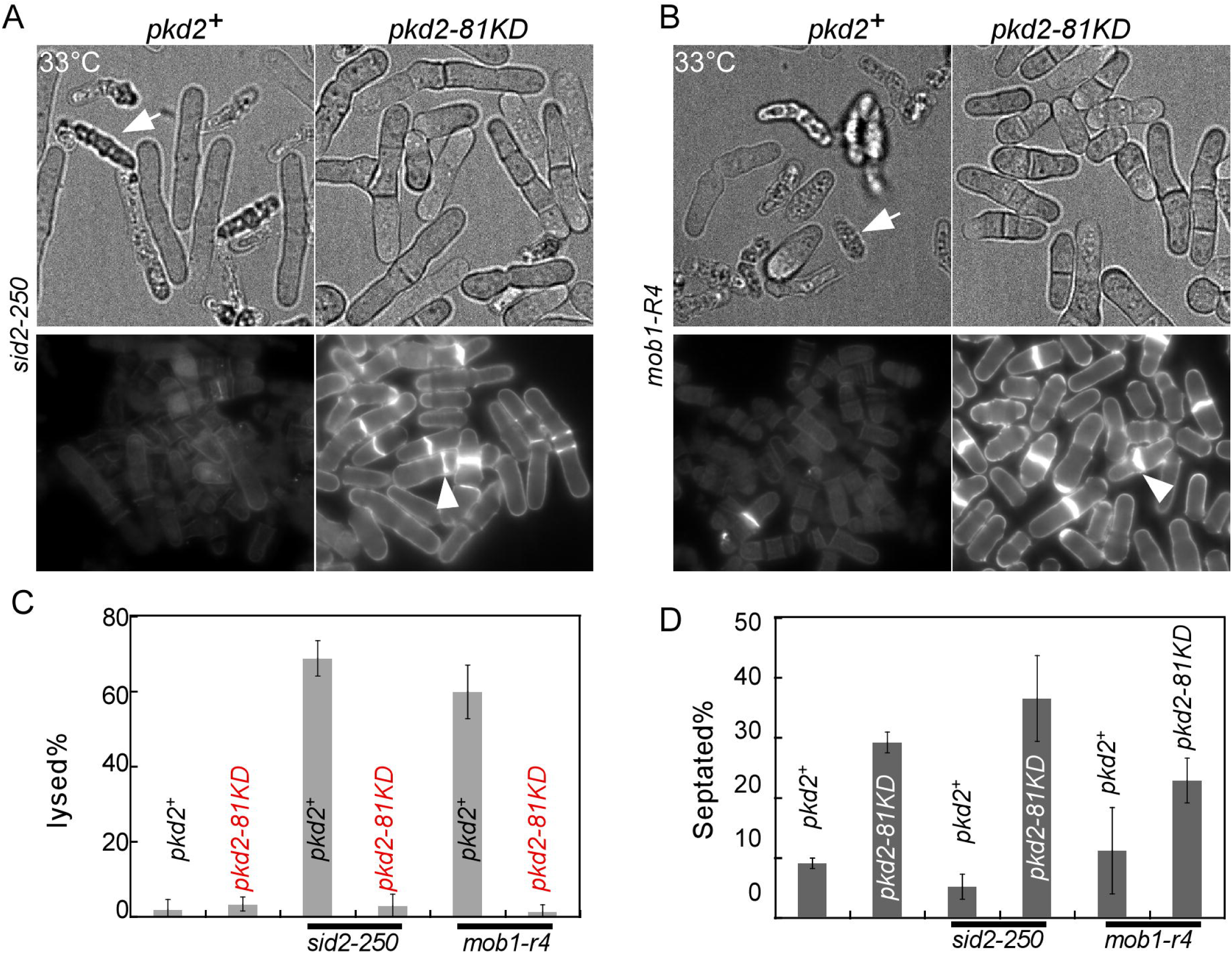
The *pkd2-81KD* mutation rescued two SIN mutants at the semi-restrictive temperature, related Fig. 7. (A-B) Micrographs of *sid2-250, sid2-250 pkd2-81KD, mob1-R4* and *mob1-R4 pkd2-81KD* cells at 33°C. Top: bright-field micrographs of the live cells. Bottom: fluorescence micrographs of the calcofluor stained cells. (C-D) Bar graphs comparing the percentage of either lysed (C) or septated (D) cells among the six strains at 33°C (n > 500). The SIN mutants frequently lysed and contained few septum at the semi-restrictive temperature, defects that were rescued by the *pkd2* mutation.

## Supplemental materials

**Movie S1: Pkd2p-GFP localized to the cleavage furrow**. Time-lapse microscopy of cells expressing Pkd2p-GFP (green) and Rlc1p-tdTomato (red). Pkd2p localized to the cleavage furrow following the contractile ring assembly.

**Movie S2: Depletion of Pkd2p by the *pkd2-81KD* mutation.** Time-lapse microscopy of *pkd2::P81nmt1-pkd2-GFP* cells in YE5s. Left: bright-field images. Right: fluorescence images. Pkd2p-GFP fluorescence was greatly reduced because of the *pkd2-81KD* mutation and many mutant cells shrunk temporarily, visible in the bright-field video.

**Movie S3: Temporary shrinking of *pkd2-81KD* cells.** Time-lapse microscopy of *pkd2-81KD* cells in YE5s. Many mutant cells deflated temporarily, visible in the bright-field video.

**Movie S4: A “wavy” contractile ring in a deflated *pkd2-81KD* cell.** Time-lapse microscopy of a *pkd2-81KD* mutant cell expressing Rlc1p-tdTomato. Temporary shrinking of the cell converted the circular ring into a “wavy band”.

**Movie S5: The microtubule turnover in wild-type cells.** Time-lapse microscopy of wild-type cells expressing Atb2p-mCherry, a marker for microtubules. Left: bright-field images. Right: fluorescence images. The interphase microtubule bundles of these wild-type cells underwent continuous turnover.

**Movie S6: Deflation of the *pkd2* mutant cells stopped the microtubule turnover.** Time-lapse microscopy of *pkd2-81KD* cells expressing Atb2p-mCherry. Left: bright-field images. Right: fluorescence images. Compared to the wild-type cells (Movie S5), turnover of the interphase microtubule bundles, stopped temporarily in the deflate mutant cells (arrows).

**Movie S7: Deflation of the *pkd2* mutant cells triggered nuclear localization of Sty1p.** Time-lapse microscopy of *pkd2-81KD* cells expressing Sty1p-GFP. Left: bright-field images. Right: fluorescence images. Nuclear localization of Sty1p-GFP increased significantly in those deflated mutant cells.

**Movie S8: The contractile ring assembly, maturation and closure in the wild-type cells.** Time-lapse fluorescence microscopy of wild-type cells expressing Rlc1p-tdTomato (Red) and Sad1-GFP (Green).

**Movie S9: The contractile ring assembly, maturation and closure in *pkd2-81KD* cells.** Time-lapse fluorescence microscopy of *pkd2-81KD* cells expressing Rlc1p-tdTomato (red) and Sad1-GFP (green). The mutant cells assembled and matured their rings more slowly but their ring closure was more rapid, compared to the wild-type cells (Movie S8). In the mutant cells, the contractile ring placement was not altered by the *pkd2* mutation and the rings didn’t slide either.

**Table S1:**
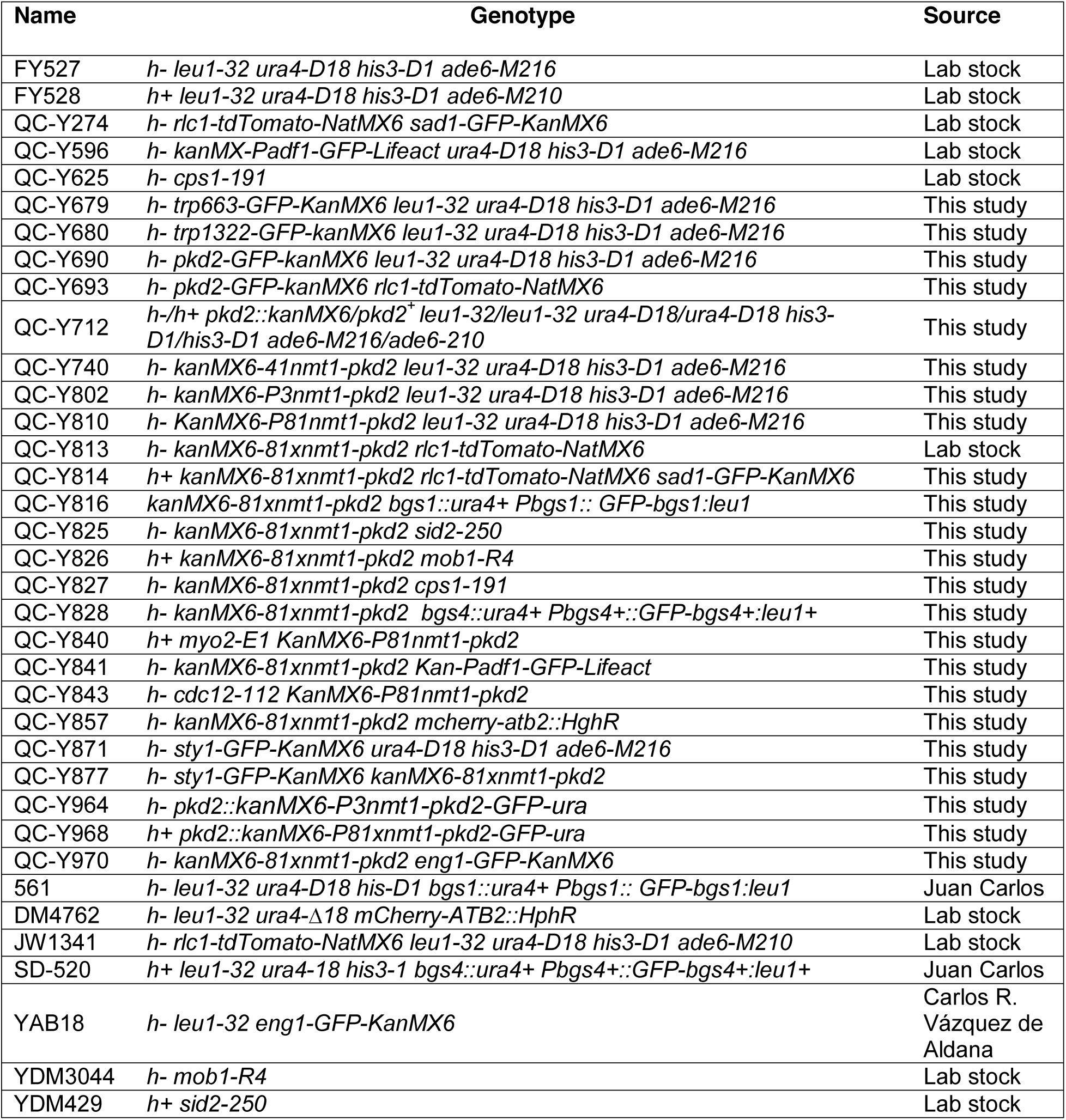
Yeast strains used in this study

